# Genomic insights into adaptation strategies and microevolutionary forces of novel non-AOA *Nitrososphaeria* in acid mine drainage ecosystems

**DOI:** 10.1101/2025.10.01.679748

**Authors:** Licao Chang, Xikai Su, Wenzhe Hu, Yun Fang, Jun Liu, Jintian Li, Linan Huang, Wensheng Shu

## Abstract

The class *Nitrososphaeria* is best known for ammonia-oxidizing archaea (AOA), yet deeply branching non-AOA lineages remain poorly characterized, leaving a critical gap in our understanding of the group’s early evolution and ecological diversification. Herein, we recovered 44 non-AOA *Nitrososphaeria* metagenome-assembled genomes (MAGs) from acid mine drainage (AMD) sediments in diverse metal mines, representing two novel genera within the family UBA164, *Acidarchaeum* and *Thermosulfuris*. A meta-analysis of 251 AMD-associated metagenomes showed that these potentially thermophilic lineages are globally distributed but typically rare, with local peaks (∼6.6%) at sites such as Fankou. Metabolic reconstruction suggested a facultatively anaerobic, mixotrophic lifestyle capable of CO oxidation and sulfur reduction, and extensive acid- and heavy-metal resistance mediated primarily by ether-linked archaeal lipids, ion efflux systems, and enzymatic reduction. Genus-specific traits include dissimilatory sulfate reduction in *Thermosulfuris* and urea utilization in *Acidarchaeum*, illuminating distinct ecological niches for them. Population-genomic analyses reveal low homologous recombination and pervasive purifying selection in these non-AOA populations, together with local relaxation of selection and elevated diversity, the former being correlated with geochemical stressors (notably copper), pointing to long-term, geochemically driven adaptation. Overall, these findings provide insights into the biodiversity, ecophysiology, and evolutionary dynamics of non-AOA *Nitrososphaeria*.

**IMPORTANCE:** Members of the class *Nitrososphaeria* that oxidize ammonia are central to the global nitrogen cycle, yet deep-branching lineages that lack this metabolism remain poorly explored, obscuring their early evolutionary trajectory. Here, we identify two new genera of non-ammonia-oxidizing *Nitrososphaeria* from acid mine drainage ecosystems and show that they recur globally in these acidic, metal-rich, oligotrophic habitats. Genomic analyses reveal adaptations for survival in harsh environments and genus-specific metabolic traits, suggesting distinct ecological strategies. Evolutionary analyses further indicate that these lineages are highly specialized, with population-genomic patterns consistent with long-term adaptation to geochemical conditions. This work sheds light on the biodiversity and evolutionary constraints of early-diverging lineages of *Nitrososphaeria*.

## INTRODUCTION

The class *Nitrososphaeria* (formerly *Thaumarchaeota*) encompasses archaea that play a fundamental role in the global nitrogen cycle, most notably the ammonia-oxidizing archaea (AOA), which catalyze the rate-limiting step of nitrification (1). Yet several deeply branching lineages within this class lack the canonical ammonia-oxidation machinery (2), implying that this metabolically defining trait was acquired later. This hypothesis is supported by a recent study that inferred this acquisition event occurred in the last common ancestor of AOA during or after the Great Oxygenation Event around 2.3 billion years ago (3). These non-AOA lineages thus provide a valuable window into the early metabolic diversification and evolutionary trajectory of *Nitrososphaeria*.

Numerous non-AOA *Nitrososphaeria* have been detected across a wide range of habitats, including marine environments (4), hot springs (5), acidic soils (6), and aquifer sediments (7), but remain poorly characterized. To date, only a single non-AOA strain, *Conexivisphaera calidus* NAS-02, has been isolated from acidic hot springs; it is a strictly anaerobic, thermotolerant, and acidophilic archaeon capable of sulfur and iron reduction (8). Comparative genomic and phylogenomic studies indicate a complex evolutionary history in *Nitrososphaeria*, including transitions from non-AOA to AOA lineages and shifts between anaerobic and aerobic respiration, driven by lateral gene transfer (LGT), gene loss, and gene duplication (3, 9–11). Despite emerging broad taxonomic patterns, fine-scale microdiversity and population-level evolutionary processes within non-AOA *Nitrososphaeria* groups are largely unexplored.

Acid mine drainage ecosystems are characterized by low pH and elevated concentrations of heavy metals and sulfate (12). Because they mimic early-Earth conditions (13), AMD ecosystems serve as natural laboratories for studying microbial adaptation to extreme selective pressures and offer important insights into ancient metabolic and evolutionary processes. Recent metagenomic surveys have revealed numerous previously undescribed archaeal lineages in AMD habitats, including non-AOA members of *Nitrososphaeria* (11). Importantly, studies show that pH is a key driver of niche specialization and evolutionary diversification in *Nitrososphaeria* (11, 14, 15). Given the strong acidity of AMD environments, their non-AOA *Nitrososphaeria* may represent an evolutionary link between early-diverging lineages and modern AOA. Consequently, AMD ecosystems provide a unique opportunity to connect genomic potential to extreme environmental constraints and to uncover the adaptive strategies of non-AOA *Nitrososphaeria*.

Herein, we applied genome-resolved metagenomics and large-scale comparative analyses to non-AOA *Nitrososphaeria* from AMD habitats. We recovered 44 MAGs and surveyed their global distribution across 251 AMD metagenomes. Metabolic reconstruction revealed strategies that enable survival in such extreme conditions, and population-genomic analyses exposed the microevolutionary forces shaping these non-AOA archaea. Collectively, this work refines the taxonomy of understudied non-AOA *Nitrososphaeria*, highlights their metabolic versatility for adaptation to extreme environments, and clarifies their evolutionary dynamics.

## RESULTS

### Genomic discovery and phylogeny of the UBA164 family in AMD sediments

A total of 44 MAGs assigned to the family UBA164 (order *Conexivisphaerales*, class *Nitrososphaeria*) were retrieved from 17 AMD sediment samples, including 11 from a lead-zinc mine, five from two copper mines, and one from a pyrite-copper mine (Tables S1 and S2). Of these MAGs, 12 are high quality, while the others are medium quality according to current genomic evaluation standards (16). Our ANI analysis identified three species-level representative genomes (FK_Bin1, FK_Bin2, and FK_Bin3) based on a 95% similarity threshold, all exhibiting high completeness (93.09 ± 2.23%) and low contamination (0.65 ± 0.56%) (Table S2). Their extrapolated genome sizes range from 1.57 to 1.72 Mbp, and complete 16S rRNA genes (1495 bp) were identified (Table S2).

To obtain their accurate taxonomic status, a phylogenomic tree was constructed based on 53 concatenated archaeal marker genes (Figure 1a). Considering relative evolutionary divergence, the phylogeny revealed that the family UBA164 was composed of two genera, JAJZYL01 (including FK_Bin1 and FK_Bin2) and UBA164 (including FK_Bin3). Notably, the AAI values among members of the UBA164 genus ranged from 59% to 90%, falling outside the genus-level threshold of 65-95% (17). Given the phylogeny and AAI values, it may be appropriate to separate the UBA164 genus in the GTDB release 220 taxonomy into the three distinct genera (Figure 1a). Herein, we propose the genus JAJZYL01 (FK_Bin1 and FK_Bin2) as *Acidarchaeum* and the FK_Bin3 and DRTY3_bin.13 cluster as a new genus *Thermosulfuris*, under the SeqCode (18). Furthermore, the AAI, ANI, and 16S rRNA gene sequence similarity values indicate that FK_Bin1 and FK_Bin2^Ts^ belong to a newly named species *Acidarchaeum fankouense*, whereas FK_Bin3^Ts^ represent a newly named species *Thermosulfuris yongpingense*. Protologues are provided in Table S3. The above results were also supported by the 16S rRNA gene phylogeny (Figure S1).

**Figure 1.**
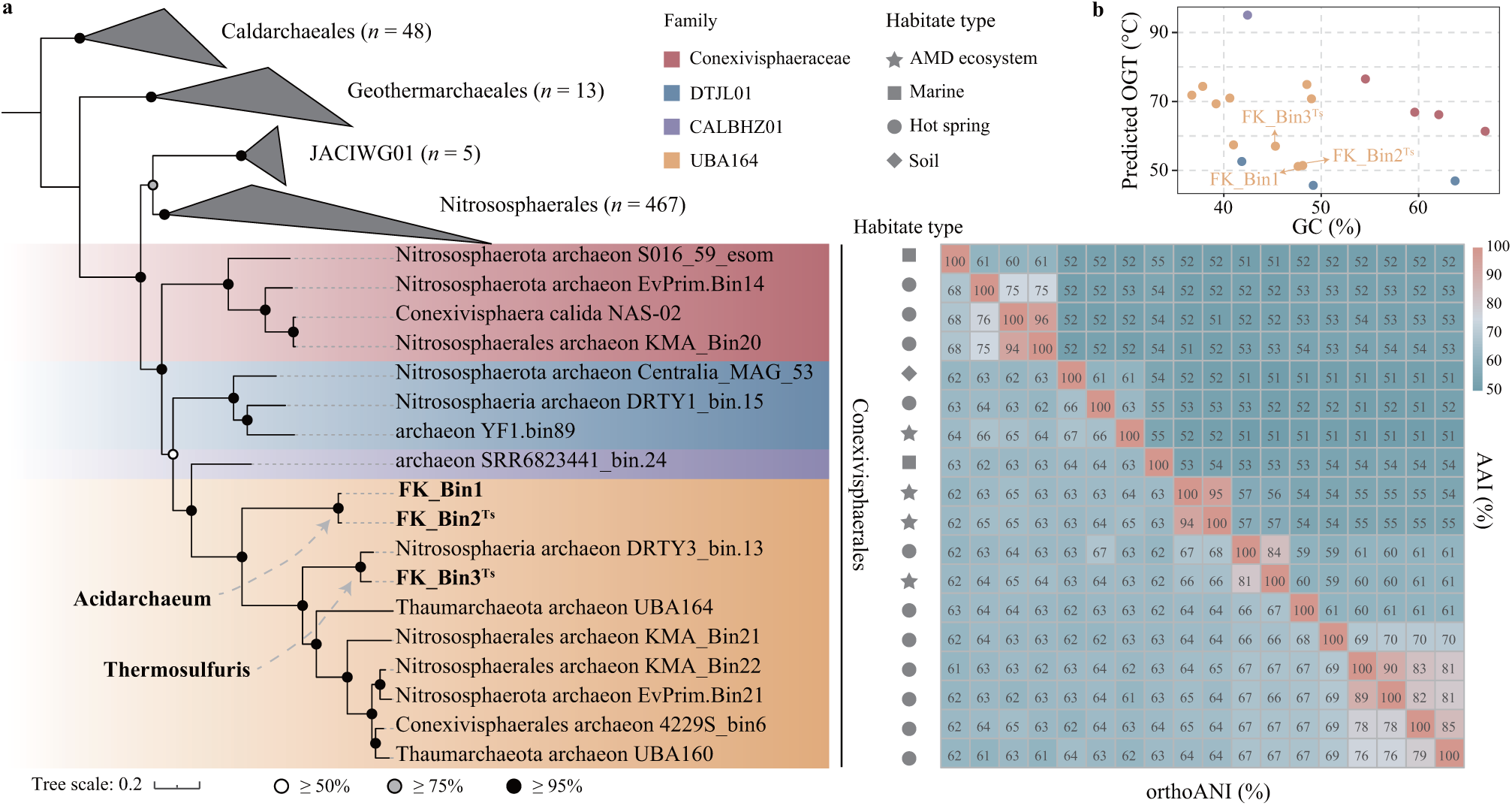
Phylogenomics, habitat distribution, and predicted optimal growth temperature of the UBA164. (a) Phylogenomic tree inferred from 53 archaeal-specific conserved marker genes, with associated habitat information, AAI and OrthoANI values. (b) Optimal growth temperature of UBA164 predicted based on genomic data.

### Distribution of UBA164 members in global AMD ecosystems

Analysis of *Conexivisphaerales* 16S rRNA gene sequences from the SILVA database (v138.2) and those retrieved from genomes in the GTDB release 220 revealed that UBA164 representative species are predominantly found in hot springs and AMD ecosystems (Figure S1). Predicted optimal growth temperatures (OGT) for UBA164 members are above 50 ℃, implying these archaea are thermophiles (Figure 1b). This aligns with the temperatures of their native environments and their observed distribution patterns. However, the scarcity of UBA164 16S rRNA gene sequences limits our insight into their geographic distribution. To address this, we performed a large-scale meta-analysis of 251 metagenomic datasets from global AMD ecosystems, including 162 AMD sediments, 58 AMD samples, 18 AMD biofilms, and 13 mine tailings, with the majority (*n* = 169) originating from China (Table S4). The results revealed that UBA164 species are broadly distributed (*n* = 166, including 139 AMD sediments, 18 AMD samples, and 9 mine tailings) but typically present at low abundance (<1%; Figure 2a and Table S4), although their relative abundances in AMD sediments from the Yongping copper and Fankou lead-zinc mines (China) could exceed 1%, reaching up to ∼6.6% (Table S4). Comparative analyses showed UBA164 to be significantly enriched in Fankou relative to other sites, mainly contributed by the higher *Acidarchaeum* abundance (Figure 2b). Given the physicochemical data (Table S5), our statistical results demonstrated that in Fankou, the UBA164 abundance declined significantly with increasing sulfate concentration, and a similar pattern was observed for *Thermosulfuris* (Figure 2c). Additionally, the relative abundance of *Thermosulfuris* also correlated significantly with lead concentration (Figure S2).

**Figure 2.**
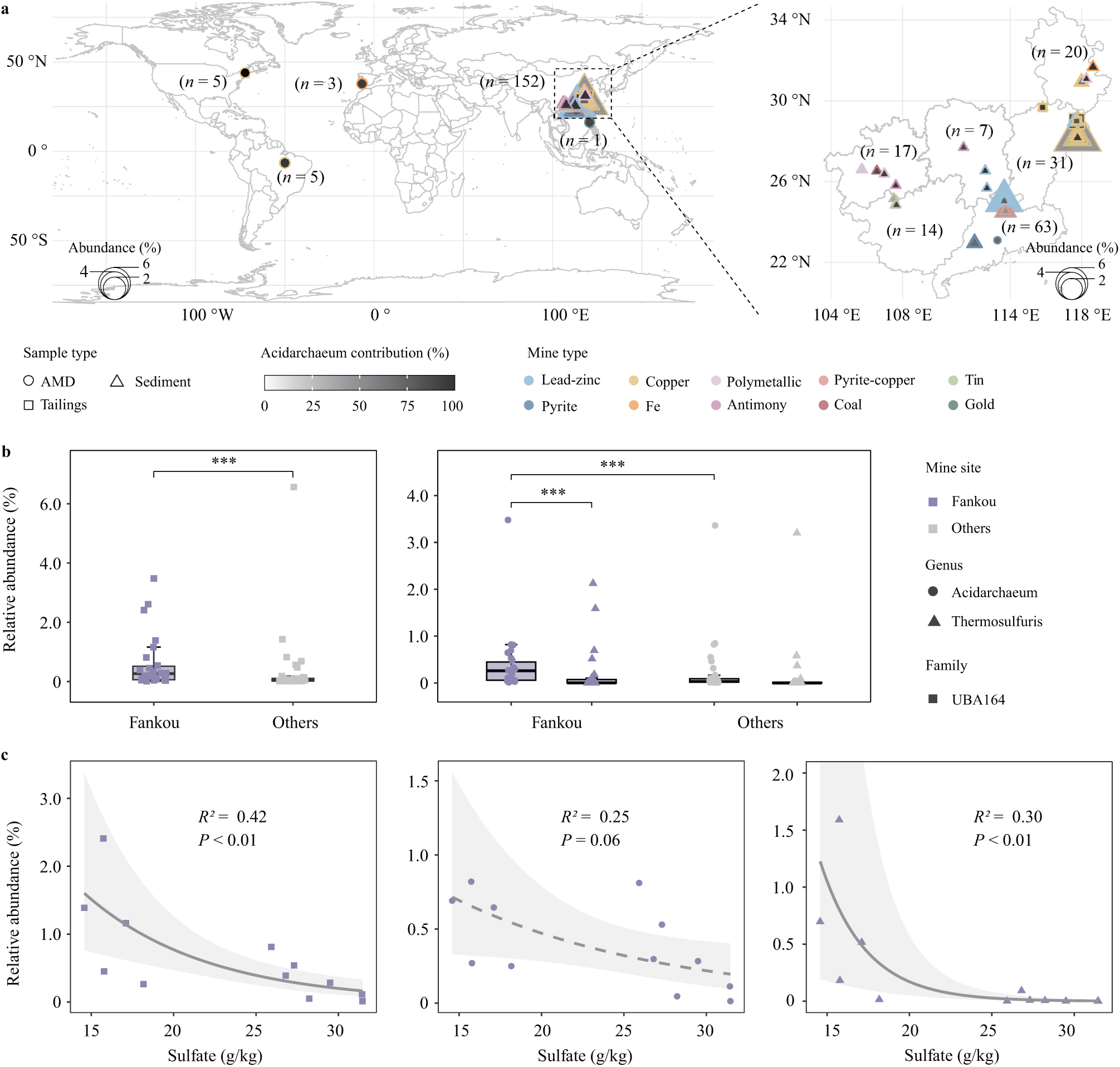
Distribution of UBA164 species in AMD environments. (a) Global distribution pattern. (b) Relative abundance of UBA164 in Fankou compared to other Chinese AMD sites. (c) Correlations between UBA164 abundance and sulfate concentration in Fankou samples.

### Metabolic potential of UBA164 species in AMD ecosystems

We reconstructed the metabolic networks of *Acidarchaeum fankouense* and *Thermosulfuris yongpingense* based on high-quality representative genomes (Figure 3 and Table S6), elucidating their ecological roles and metabolic adaptation strategies in AMD ecosystems.

**Figure 3.**
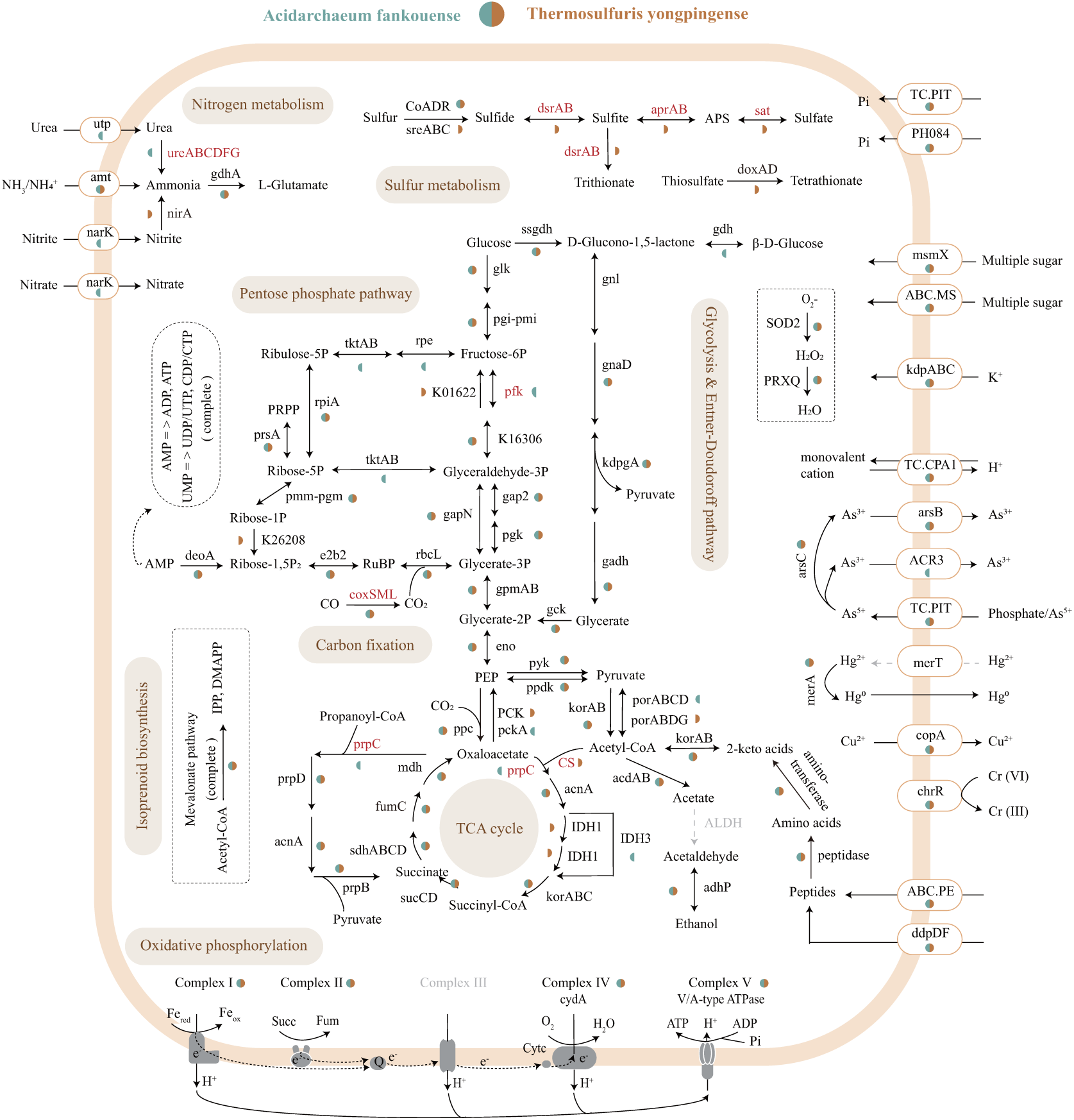
Overview of metabolic potentials of UBA164 species. Black solid arrows denote pathways present in at least one genome, gray dashed arrows indicate pathways absent in all genomes.

In the MAGs of both species, we identified the *rbcL* gene, which encodes the large subunit of ribulose-1,5-bisphosphate carboxylase/oxygenase (RuBisCO), a key enzyme for carbon fixation in the Calvin-Benson-Bassham (CBB) cycle (19). Phylogenetic analysis confirmed that all these RuBisCOs belong to the classical archaeal Form III-b (Figure S3), supporting the potential for carbon fixation in these microbes (20, 21). However, the absence of the *prk* gene (encoding phosphoribulokinase) in both species likely blocks the direct conversion of ribulose-5-phosphate to ribulose-1,5- bisphosphate. Surprisingly, the AMP nucleotide salvage pathway composed of AMP phosphorylase (*deoA*) and ribose 1,5-bisphosphate isomerase (*e2b2*) was identified, suggesting a potential role in supplying ribulose 1,5-bisphosphate for RuBisCO (22). This autotrophic capability likely provides both species with a competitive advantage in nutrient-poor environments, such as AMD sediments and mine tailings. Anaplerotic CO_2_ assimilation has also been widely reported in microbes from floodplain sediments and seawater (4, 23). Furthermore, *coxSML* genes encoding the aerobic carbon monoxide dehydrogenase (CODH) complex were detected in both species. Notably, the *cox* complex is prevalent in *Conexivisphaerales*, with all detected CoxL sequences exclusively classified as Form II (Figure 4), aligning with prior findings (11). Phylogenetic analysis revealed that these Form II CoxL sequences from the UBA164 family carry the variant motif PYRGAGR (Figure 4). Given the higher CO affinity of Form II CODH enzymes relative to Form I (24, 25), their prevalence in *Conexivisphaerales* may reflect an adaptation to environments with limited CO availability. Besides, the *coxL* gene clusters from *Conexivisphaerales* were phylogenetically placed within the *Actinomycetota* clade, suggesting they were likely acquired via HGT from *Actinomycetota*. Additionally, the presence of the *cydA* gene, encoding subunit I of the high-affinity cytochrome *bd* oxidase, hints at the possibility of a microaerophilic lifestyle, as CydA retains nearly all conserved residues essential for oxygen reduction and proton translocation (26, 27). This result corroborates recent findings that *cydA* is present exclusively in non-AOA *Nitrososphaeria* (3, 11). These species were presumed to use oxygen as an electron acceptor during CO oxidation.

**Figure 4.**
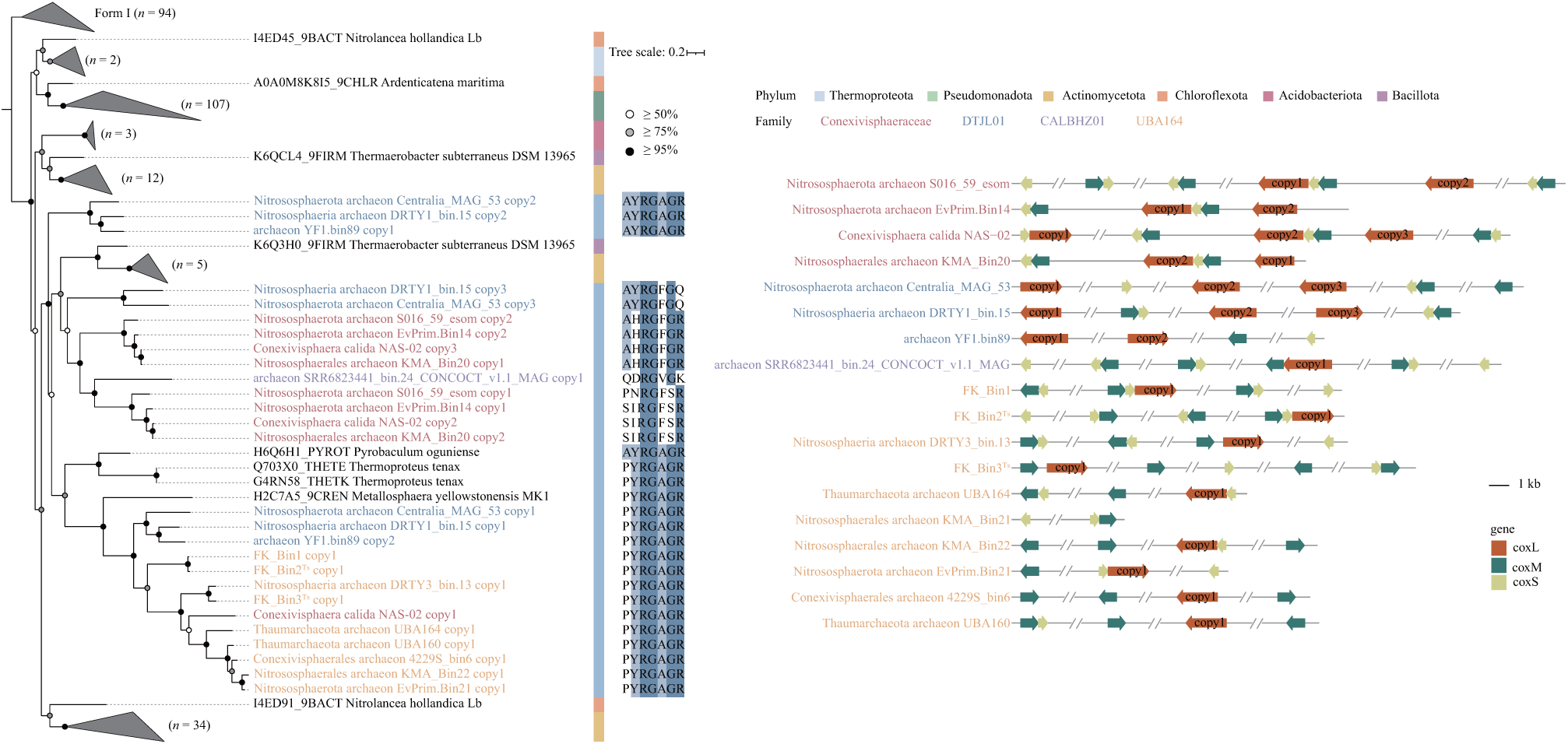
Phylogenetic analysis and operon structure of CoxL. CoxL protein sequences recovered in this study are highlighted in color, with Form I CoxL sequences used as an outgroup. Motif sequence similarity is represented by color intensity, and the operon structure of coxSML in Conexivisphaerales is shown.

Genomic analyses showed that both species likely utilize environmental peptides and proteins as carbon and energy sources. They encode peptide transporters and diverse peptidases (Tables S6 and S7) that hydrolyze oligopeptides into amino acids. Those amino acids would then be converted into keto-acids by the *aspB*- and *argD*-encoded transaminases (28), and subsequently oxidized to acetyl-CoA by 2-oxoglutarate/2-oxoacid ferredoxin oxidoreductase (*kor*), enabling their entry into central carbon metabolism (29). This process not only supplies carbon and energy but also regenerates reducing equivalents, underscoring amino acid turnover as an internal energy reserve in sediment-dwelling archaea (30). Such reliance on exogenous peptides and proteins is likely a common strategy among archaea inhabiting extreme environments, such as AMD and acidic hot springs, where detrital inputs are abundant (31).

Dissimilatory sulfate/sulfite reduction, among the earliest known microbial metabolic processes (32), also serves as an important energy source for anaerobic microorganisms (33). In *Thermosulfuris yongpingense* FK_Bin3^Ts^, the complete pathway composed of sulfate adenylyl transferase (*sat*), adenosine 5’-phosphosulfate reductase (*aprAB*), and dissimilatory sulfite reductase (*dsrAB*) was identified (Figure 3). The absence of *dsrEFH* and *dsrL* genes in its *dsr* operon, along with the phylogenetic placement of DsrAB, supports the potential for dissimilatory sulfate reduction in this archaeon (Figure S4a) (34, 35). Moreover, *dsrAB* genes are unevenly distributed across the phylum *Thermoproteota*. In the classes *Thermoprotei* and *Nitrososphaeria*, the encoded DsrAB proteins are of the reduced archaea type, whereas those in *Korarchaeia* and *Nitrososphaeria_A* belong to the reduced bacteria type (Figure S4) (36). This pronounced phylogenetic divergence suggests that *dsrAB* genes were not vertically inherited from a common ancestor. Instead, their patchy distribution likely reflects multiple independent HGT events, implying that the last common ancestor of *Thermoproteota* may have entirely lacked *dsrAB*. Furthermore, within the order *Conexivisphaerales*, *dsrAB* genes are found exclusively in *Thermosulfuris* and a neighboring genus, both members of the family UBA164 (Figure S4). Their phylogenetic clustering with sequences from *Thermoprotei* suggests HGT from *Thermoprotei* to this genus. Given the high sulfate concentrations and near-zero oxygen in AMD ecosystems and acidic hot springs (37), *Thermosulfuris* members (FK Bin3^Ts^ and DRTY3_bin.13) likely use sulfate/sulfite, rather than oxygen, as the electron acceptor for CO oxidation (25). In addition, both *Acidarchaeum fankouense* and *Thermosulfuris yongpingense* exhibit genetic potential for S^0^ and S_2_O_3_^−^ reduction, as indicated by the presence of *sreABC* and/or *CoADR* genes, which encode sulfur reductase and CoA-dependent NAD(P)H sulfur oxidoreductase, respectively. This highlights the significant roles of these species in sulfur cycling in AMD ecosystems.

Urea, a prevalent dissolved organic nitrogen, is a readily available nitrogen source for aquatic microorganisms because it can be rapidly taken up and hydrolyzed (38, 39). The complete urea utilization pathway, including the urea transporter (*utp*) and the urease operon (*ureABC*), was found exclusively in *Acidarchaeum fankouense* (Figure 3). This pathway is also essential for many AMD-inhabiting microbes as an important nitrogen source (37, 40). Given that urease was absent in the last common ancestor (LCA) of *Nitrososphaeria* (23), this species likely acquired the urea utilization ability via HGT during evolution. Besides, the *nirA* gene, encoding ferredoxin-nitrite reductase, was identified in *Thermosulfuris yongpingense* FK_bin3^Ts^, implying its ability to assimilate nitrite as a nitrogen resource. For inorganic phosphorus utilization, both species can hydrolyze inorganic pyrophosphate (PPi) to orthophosphate via pyrophosphatase (*ppa*) (Table S6), harnessing the energy released to support growth (41). In brief, these UBA164 archaea inhabiting AMD ecosystems are facultative anaerobic mixotrophic CO-oxidizers with distinct roles in biogeochemical cycling.

### Environmental adaptation

AMD ecosystems, characterized by low pH and high concentrations of toxic metals (42, 43), exert strong selective pressure on microorganisms, driving the evolution of specialized adaptive mechanisms (40, 44). To cope with acid stress, *Acidarchaeum fankouense* and *Thermosulfuris yongpingense* employ multiple strategies. Results showed that both species possess a complete, modified mevalonate (MVA) pathway for isoprenoid biosynthesis, supported by the presence of key genes encoding mevalonate 5-phosphate dehydratase (*acnx1* and *acnx2*) and anhydromevalonate phosphate decarboxylase (Tables S6). Isoprenoid-derived ether bonds are known to reduce membrane permeability to small molecules, thereby enhancing archaeal stability under acidic conditions (45). This variant of the MVA pathway, first identified in *Aeropyrum pernix* and considered to represent the most ancient form, requires less ATP. This energy-efficient trait confers a survival advantage in energy-limited anaerobic conditions and likely supports the survival of UBA164 members in AMD environments (46). Moreover, both species likely maintain a near-neutral cytoplasmic pH through the activity of the high-affinity potassium transport system (KdpABC), Na^+^:H^+^ antiporters, and cytoplasmic buffer molecules such as glutamate, arginine, and lysine (47, 48). These acid resistance strategies are commonly employed by microbes inhabiting AMD ecosystems (40, 44).

To counter heavy metal stress, both species likely rely primarily on coordinated ion efflux and enzymatic reduction. For instance, the *arsBCR* operon enables the reduction of As(V) to As(III) via ArsC, followed by export through the ArsB pump (49). They also likely encode the copper transporter CopA for Cu^2+^ efflux, a common defense among AMD microbes (50). Additionally, these UBA164 archaea appear capable of reducing Cr(VI) to the less-toxic Cr(III) via ChrR and Hg²⁺ to volatile Hg⁰ through MerA (51, 52). The presence of a vacuolar iron transporter (VIT) further suggests a strategy for sequestering excess Fe²⁺, helping to mitigate cytotoxicity and maintain iron homeostasis (53). In response to oxidative stress, both species encode a variety of defense proteins, including peroxiredoxin 2/4 (*ahpC*), superoxide dismutase (SOD2), and thioredoxin-dependent peroxiredoxin (PRXQ), consistent with observations in other AMD-associated microbes (54). They also produce three molecular chaperones, GrpE, DnaJ, and DnaK, that protect intracellular proteins and nucleic acids from oxidative damage. This protective mechanism is evident in *Leptospirillum ferriphilum* inhabiting AMD biofilms, where protein-refolding chaperones are highly expressed (55). In short, these findings illuminate the stress resistance mechanisms that enable UBA164 archaea to survive and thrive in AMD ecosystems.

### Genomic variations across UBA164 species in AMD ecosystems

To better understand the microevolution of UBA164 species in AMD environments, we calculated diverse evolutionary metrics using representative MAGs of *Acidarchaeum fankouense* and *Thermosulfuris yongpingense*, including linkage disequilibrium (D′), nonsynonymous to synonymous mutation ratio (pN/pS), and nucleotide diversity (SNVs/kbp). High D′ values observed in Fankou (0.97 ± 0.02) and other sites (0.98 ± 0.03) suggest low levels of homologous recombination in UBA164 populations (Figure 5a). This finding also implies that these populations may inhabit relatively constant environments where genetic variation likely arises through mutation and/or drift (56). This phenomenon is prevalent in nature, such as sulfate-reducing microbes in a deep-sea cold seep sediment (57). Moreover, all pN/pS ratios for UBA164 populations were well below one, although significantly higher values were observed in Fankou (Figure 5a). This pattern indicates that UBA164 populations in Fankou experience more relaxed purifying selection than those in other sites, despite strong purifying selection being maintained across all sites. Interestingly, the pN/pS values of UBA164 were negatively correlated with copper concentrations in Fankou (Figure S5). This suggests that elevated copper levels impose toxic stress that strengthens purifying selection on these archaea, consistent with prior reports of metal-driven selection (58). Given the significantly higher intra-population diversity (SNVs/kbp) and pN values in Fankou, and no corresponding difference in pS values across sites (Figure 5a), UBA164 microdiversity in Fankou appears to be primarily driven by the accumulation of nonsynonymous mutations. Nevertheless, across nearly all ecosystems, microbial populations experience purifying selection, with the majority of nucleotide variation remaining neutral (59, 60). Taken together, these results imply that these UBA164 populations are highly adapted to the relatively stable AMD environments, especially Fankou.

**Figure 5.**
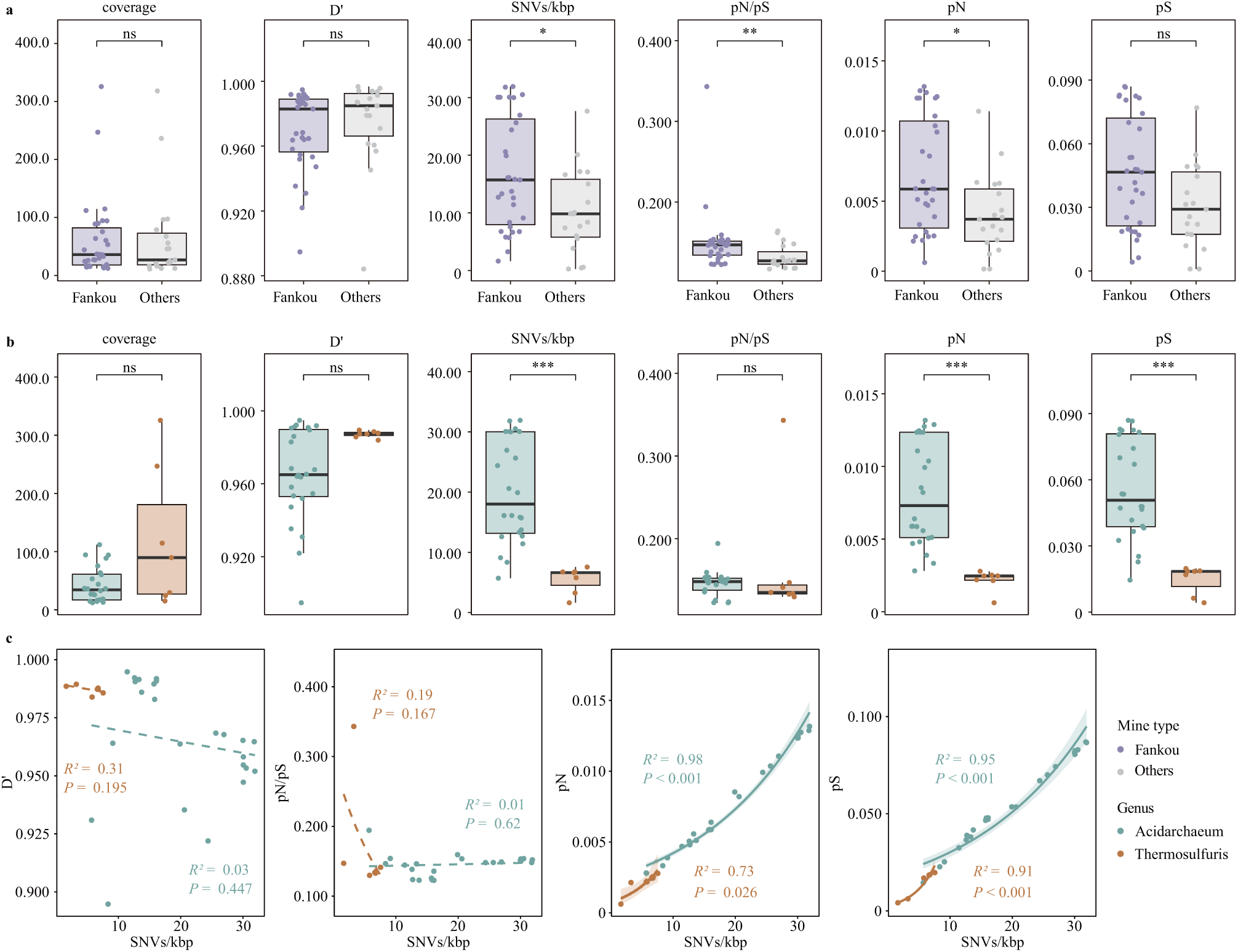
Microdiversity patterns of UBA164 species. (a) Microdiversity comparison between Fankou and other Chinese AMD sites. (b) Microdiversity comparison between Acidarchaeum and Thermosulfuris in Fankou. (c) Correlation of evolutionary metrics for UBA164 species in Fankou.

In the Fankou samples, the genera *Acidarchaeum* and *Thermosulfuris* displayed distinct evolutionary trajectories despite comparable levels of purifying selection and recombination. Specifically, *Acidarchaeum* populations harbored significantly greater standing genetic variation than *Thermosulfuris*, characterized by proportional increases in both pS and pN (Figure 5b). This pattern is more consistent with differences in long-term demographic history rather than a reduction in the intensity of purifying selection (61). Taken together with their functional differences, these results indicate that microbial populations occupying distinct niches in AMD environments adopt diverse evolutionary strategies, a pattern also observed in deep-sea hydrothermal vents and cold seep sediments (57, 62). Within each genus, pN and pS increase with SNV density, whereas pN/pS and D′ do not (Figure 5c), indicating that rising microdiversity reflects an expansion of the polymorphic site pool under mutation-selection-drift balance rather than changes in selective constraint or recombination rate (63, 64). In addition, *Acidarchaeum* populations showed a significant positive correlation between SNVs/kbp and genome coverage (Figure S6), indicating that larger populations harbor greater genetic variation associated with an expanded polymorphic site pool that underpins future adaptive potential (57). Overall, these results demonstrate that purifying selection is the primary force shaping UBA164 populations in AMD environments, with its strength modulated by local geochemistry, such as copper stress. Furthermore, co-occurring UBA164 lineages, namely *Acidarchaeum* and *Thermosulfuris*, employ distinct evolutionary strategies, a pattern also observed in *Sulfurovum* species from the geochemically contrasting Piccard and Von Damm vent fields (62).

## DISCUSSION

Ammonia-oxidizing *Nitrososphaeria* are pivotal players in the nitrogen cycle (1), yet deeply branching, non-AOA lineages remain poorly characterized. Here, we recovered three high-quality UBA164 MAGs from AMD habitats, environments that recapitulate aspects of early-Earth geochemistry (13), and described two novel genera, *Acidarchaeum* and *Thermosulfuris*. These genomes expand the known phylogenetic breadth of non-AOA *Nitrososphaeria*, and provide an opportunity to disentangle their distribution, adaptive mechanisms, ecological roles, and evolutionary history.

Although these UBA164 lineages are globally detectable in AMD sediments, they are typically rare and only occasionally bloom locally (e.g., Fankou). Their predicted thermophily aligns with proposals of a hot origin for *Nitrososphaeria* (11), a trait that is unusual in predominantly mesophilic AMD communities (12, 65, 66). We hypothesize that their sporadic presence reflects colonization within transient, localized hot spots generated by exothermic iron-sulfur mineral oxidation (67); this model helps to reconcile both their overall rarity and the compositional similarity sometimes observed between AMD and acidic hot-spring communities (65, 68).

Metabolic inference reveals considerable versatility. These archaea encode Form II CODHs, cytochrome bd oxidase, and Form III-b RuBisCO paired with an AMP-salvage route, alongside extensive peptidolytic and transport systems. Collectively, these pathways indicate facultative anaerobic, mixotrophic lifestyles in which CO oxidation can supply energy across a range of redox conditions and heterotrophic peptide degradation supports organic-carbon assimilation (11). *Thermosulfuris* additionally encodes a complete dissimilatory sulfate reduction pathway, suggesting sulfate can serve as an alternative terminal electron acceptor under anoxic conditions; *Acidarchaeum* encodes urease, pointing to an auxiliary nitrogen source. This complementarity, together with the capacity to operate as both autotrophic carboxydotrophs and heterotrophic carboxydovores, using either oxygen or sulfate as terminal electron acceptors (25), supports a model of niche partitioning that likely reduces competition and stabilizes communities in chemically heterogeneous, nutrient-poor AMD sediments.

The patchy phylogenetic distribution of key metabolic genes, notably *coxL* and *dsrAB*, implicates HGT as an important driver shaping the metabolic repertoires of UBA164. Instances where *coxL* clusters with *Actinomycetota*-like sequences and the inferred acquisition of *dsrAB* from *Thermoprotei* illustrate how HGT can rapidly expand metabolic capacity, allowing recipients to exploit novel energy pathways in extreme environments (69, 70). Such gene flow complicates reconstruction of ancestral traits within *Nitrososphaeria* but highlights the evolutionary plasticity that supports survival under strong selective pressures (69, 71).

Population-genomic analyses reveal low homologous recombination and pervasive purifying selection across these UBA164 populations, consistent with long-term specialization to stable, harsh AMD niches (72, 73). Notably, Fankou populations exhibit elevated microdiversity and relaxed purifying selection, the latter of which correlates with copper concentrations, indicating that local geochemistry may modulate evolutionary dynamics, potentially promoting diversification through relaxed constraint or episodic selection (60, 73, 74). Distinct microevolutionary patterns between *Acidarchaeum* and *Thermosulfuris* (e.g., higher standing genetic variation in *Acidarchaeum* and different coverage-SNV relationships) suggest lineage-specific demography or ecological strategies. In summary, these UBA164 archaea are metabolically flexible, stress-tolerant lineages whose ecology and evolution are closely linked to AMD geochemistry. Given AMD’s analogy to certain early-Earth settings, these lineages offer a valuable model for exploring pre-AOA *Nitrososphaeria* physiology and the evolutionary routes that ultimately led to ammonia oxidation.

Overall, this study recovered three non-AOA *Nitrososphaeria* MAGs from AMD ecosystems, identifying two novel genera, *Acidarchaeum* and *Thermosulfuris*, within the family UBA164. These archaea are globally widespread in AMD environments but typically rare, with abundance peaks in specific sediments. Metabolic reconstructions suggest a facultative anaerobic, mixotrophic lifestyle, with genus-specific adaptations indicating distinct ecological niches. Both genera possess extensive genetic mechanisms for tolerating extreme acidity and heavy metals. Population-genomic analyses revealed that these UBA164 archaea are highly adapted to their niches, undergoing low homologous recombination and pervasive purifying selection. Site-specific genomic diversity (notably higher diversity and relaxed selection in Fankou), linked to local geochemistry like copper concentration, suggests long-term adaptation to relatively stable and geochemically defined habitats. Due to metagenomic inference and a strong geographic sampling bias in this study, future work should expand genomic sampling across diverse geographies, pursue cultivation for physiological validation, and employ *in situ* and omics analyses to confirm metabolic inferences and define ecological roles. Together, these efforts will deepen understanding of non-AOA *Nitrososphaeria* ecology, evolution, and their roles in acidic, metal-rich ecosystems.

## MATERIALS AND METHODS

### Dataset acquisition

A total of 34 samples from AMD environmental, including AMD and sediment, were collected from six different sites between 2016 and 2018 (Table S4). Sampling, DNA extraction, metagenomic sequencing, and geochemical analyses were performed based on established methods (54). Briefly, approximately 50 L of acidic water and 10 g of sediment per sample were used for DNA extraction. The extracted DNA was purified and sequenced on Illumina HiSeq or MiSeq platforms. pH was measured using a pH meter, and sulfate concentrations were determined using the BaSO4-based turbidimetric method (75). Additionally, 217 publicly available metagenomic datasets associated with AMD environments, including samples of AMD, sediment, biofilm, and tailings from ten countries, were obtained from the NCBI SRA database (https://www.ncbi.nlm.nih.gov/) and the NODE database (https://www.biosino.org/node/home). Detailed information is provided in Table S4.

### Metagenome assembly and genome binning

Metagenomic raw reads from 17 sediment samples were processed to remove duplicates using an in-house Perl script, followed by adapter removal with BBduk (Table S1) (76). Low-quality reads were filtered using Sickle v1.33 with the parameters “-q 30 -l 50” (77). Quality-controlled reads from each sample were individually assembled using SPAdes v3.15.5 with the parameters “--meta -k 33,55,77,99”. The resulting scaffolds were binned using MetaBAT2 v2.12.1 (78), MaxBin2 v2.2.6 (79), and CONCOCT v1.0.0 (80). These initial bins were consolidated using the bin_refinement module of metaWRAP v1.3.2 with parameters “-c 50 -x 10” (81), and their preliminary taxonomic classification were inferred with GTDB-Tk v2.4.0 (82). All MAGs classified under the UBA164 lineage were selected for further analysis. To improve genome quality, high-quality reads from these 17 samples were mapped to a combined UBA 164 genome dataset, which included MAGs recovered in this study, previously published UBA164 genomes, and GTDB release 220 representative genomes, using BBMap with a minimum alignment identity of 95%. The recruited reads were co-assembled and binned following the same pipeline as described above (11, 83). All obtained UBA164 MAGs were then dereplicated with dRep at 95% ANI to obtain representative genomes at the species level (84, 85). Genome quality metrics, including completeness, contamination, and strain heterogeneity, were assessed using CheckM v1.2.2 in “lineage workflow” mode. Additionally, SSU rRNA genes were identified extracted from genomes using the “ssu_finder” command implemented in CheckM (86). This analysis ultimately yielded three high-quality genomes that surpassed the quality of both currently available GTDB representatives and other previously reported UBA164 genomes (11).

### Functional annotation and pathway construction

Protein-coding sequences (CDSs) in the UBA164 genomes were predicted using Prodigal v2.6.3 with the “-p single” option (87). The predicted CDSs were then functionally annotated against the Kyoto Encyclopedia of Genes and Genomes (KEGG) database using BlastKOALA, and against EggNOG and NCBI-nr databases using DIAMOND with an E-value cutoff of 1e-5 (88, 89). Subcellular localization of predicted proteins was predicted with PSORTb v3.0.3 (90). The optimal growth temperature for each genome was inferred by Tome (91).

### Ecological distribution

To investigate the global distribution and quantify the abundance of the three representative MAGs recovered in this study, high-quality reads from 251 globally distributed metagenomic samples were mapped against these MAGs using BBmap with a minimum alignment identity of 95%. The relative abundance of each MAG within a sample was calculated as the proportion of reads mapping to it relative to the total reads in that library. A MAG was considered present in a sample if at least 20 reads were mapped to it.

### Phylogenetic analyses

A concatenated alignment of 53 archaeal marker proteins was generated by GTDB-Tk, incorporating MAGs recovered in this study along with representative genomes of the class *Nitrososphaeria* from GTDB release 220 (82). Poorly aligned regions were trimmed using trimAL v1.4 with the parameters “-gt 0.95 -cons 50” (92). A maximum-likelihood tree was constructed with IQ-TREE2 v2.2.0.3, using the best-fit model selected by ModelFinder with the “MFP” option (93, 94). Branch support was assessed with 1000 UFBoot replicates, and a hill-climbing nearest neighbour interchange (NNI) search was performed to reduce the risk of overestimating branch supports.

For the 16S rRNA gene phylogenetic tree, reference sequences affiliated with *Nitrososphaeria* were obtained from the SILVA database 138.2 and dereplicated at 95% identity using CD-HIT v4.8.1 (95, 96). These sequences, along with the 16S rRNA gene sequences derived from UBA164 genomes recovered in this study, were aligned using the SINA aligner v1.2.12 through the SILVA web interface (97). Filtering and subsequent phylogenetic analysis were performed as described above. Additionally, habitat information for the 16S rRNA gene sequences belonging to the order *Conexivisphaerales* were compiled to characterize distribution patterns.

For the phylogenetic analysis of CoxL, DsrAB and RbcL proteins, sequence alignment was performed using MAFFT v7.515 with the “--auto” parameter (98), followed by trimming with trimAL and tree construction with IQ-TREE2 as previously described. For CoxL, initial protein sequences were obtained from the UniRef90 database by querying “Carbon monoxide dehydrogenase large chain” and applying a gene name filter for “coxL” or “cutL”. Putative Form II CoxL sequences were identified by the presence of the diagnostic motifs “AYRGAGR” or “PYRGAGR”. From these, sequences >600 amino acids with unambiguous taxonomic assignments were retained, yielding 172 sequences. A phylogenetic tree was constructed using these sequences together with 94 previously reported Form I CoxL sequences (serving as outgroup) and 27 CoxL sequences from UBA164 genomes obtained in this study (99). For DsrAB and RbcL, reference sequences were obtained from previously published studies (21, 36, 54).

### Calculation of evolutionary metrics

Quality-controlled reads from each sample were mapped to representative MAGs using Bowtie2 v2.2.5 with default parameters (100). MAGs with ≥10× coverage in a given sample were retained for calculating evolutionary metrics, including D’, SNVs/kbp, and pN/pS. Genome-wide metrics were calculated from the mapping results using the “profile” module of inStrain v1.3.9 with the parameter “--database mode” (101).

### Statistical analyses

All statistical analyses were conducted in R v4.3.2. Wilcoxon rank-sum test was used to compare (i) the abundance of different microbial populations within the same site, (ii) evolutionary metrics between the Fankou site and other sites, (iii) evolutionary metrics between two distinct microbial populations. Generalized linear models (GLMs) were used to assess the relationships between species abundance and physicochemical parameters, between evolutionary metrics and physicochemical parameters, and among evolutionary metrics themselves.

## DATA AVAILABILITY

The three MAGs retrieved in this study have been deposited in the NCBI database with the accession numbers SAMN48438030, SAMN48438034, and SAMN48438039.

## ACKNOWLEDGMENTS

This work was supported by the National Natural Science Foundation of China (Nos. 42177178, 42377114, 42577235, and 42077281) and Huazhong Agricultural University Scientific & Technological Self-innovation Foundation (No. 2662025ZHPY006).

